# Consistency of EEG source localization and connectivity estimates

**DOI:** 10.1101/071597

**Authors:** Keyvan Mahjoory, Vadim V. Nikulin, Loïc Botrel, Klaus Linkenkaer-Hansen, Marco M. Fato, Stefan Haufe

**Affiliations:** Department of Informatics, Bioengineering, Robotics and System Engineering, University of Genova, Genova, Italy; Machine Learning Department, Technische Universität Berlin, Berlin, Germany; Neurophysics Group, Charité University Medicine Berlin, Berlin, Germany; Center for Cognition and Decision Making, National Research University Higher School of Economics, Russian Federation; Institute of Psychology, University of Würzburg, Würzburg, Germany; Department of Integrative Neurophysiology, Center for Neurogenomics and Cognitive Research (CNCR), Amsterdam, The Netherlands

**Keywords:** Electroencephalography (EEG), Source Localization, Functional/Effective Connectivity, Forward/Inverse Modeling, Consistency, Reproducibility

## Abstract

As the EEG inverse problem does not have a unique solution, the sources reconstructed from EEG and their connectivity properties depend on forward and inverse modeling parameters such as the choice of an anatomical template and electrical model, prior assumptions on the sources, and further implementational details. In order to use source connectivity analysis as a reliable research tool, there is a need for stability across a wider range of standard estimation routines. Using resting state EEG recordings of N=65 participants acquired within two studies, we present the first comprehensive assessment of the consistency of EEG source localization and functional/effective connectivity metrics across two anatomical templates (ICBM152 and Colin27), three electrical models (BEM, FEM and spherical harmonics expansions), three inverse methods (WMNE, eLORETA and LCMV), and three software implementations (Brainstorm, Fieldtrip and our own toolbox). Source localizations were found to be more stable across reconstruction pipelines than subsequent estimations of functional connectivity, while effective connectivity estimates where the least consistent. All results were relatively unaffected by the choice of the electrical head model, while the choice of the inverse method and source imaging package induced a considerable variability. In particular, a relatively strong difference was found between LCMV beamformer solutions on one hand and eLORETA/WMNE distributed inverse solutions on the other hand. We also observed a gradual decrease of consistency when results are compared between studies, within individual participants, and between individual participants. In order to provide reliable findings in the face of the observed variability, additional simulations involving interacting brain sources are required. Meanwhile, we encourage verification of the obtained results using more than one source imaging procedure.

## 1. Introduction

Two major methodological challenges in noninvasive neuroimaging concern the determination of task-specific cortical areas and the determination of their interactions from functional data.

Functional magnetic resonance imaging (fMRI) measures changes in blood flow induced by neuronal activity. While being able to distinguish brain activations even a few millimeters apart, fMRI suffers from poor temporal resolution with sampling rates typically lower than 1 Hz.

Compared to fMRI, electro- and magnetoencephalography (EEG/MEG) provide much higher temporal resolution thus making them attractive techniques for studying interactions between different brain structures.

Yet, EEG and MEG suffer from low spatial resolution since only superpositions of brain signals originating from the entire cortical gray matter can be recorded. Sensor space analyses in general are not suitable to infer the involvement of brain structures in interaction even in such broad terms as ‘frontal-to-occipital’ (Haufe, 2011; Van de Steen et al., 2016). The interpretation of EEG/MEG data in neuroanatomical terms therefore requires a reconstruction of the sources from the recorded data. This, however, requires a solution of an ‘ill-posed’ inverse problem, for which infinitely many solution exists. To select a unique solution, prior knowledge of the source characteristics needs to be employed. Consequently, there is a host of methods estimating sources under specific assumptions.

The choice of an inverse method is a factor that heavily influences the reconstructed brain activity, as well as subsequent analyses relying on the recovered sources. Other important factors are the specifics of the physical model of electrical current flow in the head and the choice of an anatomical template with which to perform the source reconstruction. In practice, researchers typically resort to one of the various publicly available toolboxes for source analysis such as Brainstorm (Tadel et al., 2011), FieldTrip (Oostenveld et al., 2010), EEGLAB (Delorme and Makeig, 2004) and MNE (Gramfort et al., 2014). These toolboxes typically provide ready-made anatomical templates, methods for electrical forward calculations, and implementations of inverse solutions. While the methods portfolios provided by different toolboxes are in general similar, the different possible combinations of forward and inverse models, as well as the differences in their implementations and the choice of their numerous parameters (such as tissue conductivities, segmentation and meshing parameters for forward models, and regularization and depth weighting constants for inverse models) may lead to a substantial variability of possible source location and connectivity estimates.

Numerous studies have quantified biases in the localization of brain sources (e.g., Darvas et al., 2004; Haufe, 2011; Gramfort et al., 2013), as well as in the determination of brain connectivity (e.g., Schoffelen and Gross, 2009; Haufe et al., 2010, 2013; Ewald et al., 2013; Rodrigues and Andrade, 2015; Haufe and Ewald, 2016) for specific methods. The error of a statistical measure depends however not only on its estimation bias but also on its variance. Large variability in combination with the small sample sizes that are common in neuroimaging studies have been identified as the major cause of the lack of reproducibility that is generally observed (Button et al., 2013). A recent study by Colclough et al. (2016) consequently assessed the consistency of MEG source connectivity metrics across different datasets.

When working with EEG/MEG source estimates, another source of variability to be considered is the choice of the forward and inverse modeling parameters. Intuitively, we would consider results based on reconstructed sources only meaningful if they are reasonably consistent across a range of widely accepted estimation procedures (pipelines) when applied to the same data. An investigation of this latter factor would help to assess the reliability of EEG and MEG based brain connectivity estimation as a research tool, but has not yet been provided.

With this work, we present the first comprehensive assessment of the consistency of EEG source location and connectivity analyses across common forward and inverse models. Our data is based on reconstructions performed in three different analysis packages using combinations of three different inverse methods, three different electrical modeling approaches, and two different template anatomies. We investigated the sources and communication patterns of alpha-band (8–13 Hz) oscillations using resting-state recordings acquired within two different studies (N=65). We chose to use alpha oscillations because: 1) they have high signal-to-noise ratio – thus ameliorating the problem of noisy recordings and 2) these oscillations have relatively stable spatial patterns across subjects corresponding to sources in occipito-parietal and central areas of the cortex.

Our main goal was to bring the attention of the neuroimaging community to the problem of identifying interacting neuronal sources on the basis of the multichannel EEG recordings. We wanted to illustrate pitfalls in obtaining measures of connectivity due to different stages of the data analysis including selection of the toolbox, forward/inverse models and connectivity estimates. By making researchers aware of multiple problems in connectivity analysis, we hope to help them with the validation of the results and consequently in establishing reliable findings about the brain functioning.

The paper starts by introducing the data, preprocessing steps, forward and inverse modeling approaches, and robust connectivity measures. In the experimental part we first demonstrate that the choice of the reference electrode dramatically influences EEG sensor-space connectivity maps, making sensor-space analysis unsuitable for the study of brain connectivity. Using pairwise correlations, we quantified the similarity of inverse solutions and source connectivity matrices when different source reconstruction pipelines are applied to the same data. We also quantified the within-participant, between-participant and between-study variability. We conclude the paper with a discussion of the different sources of variability and their impact on the reliability of results, strategies to deal with variability, general validation strategies, and the perhaps counter-intuitive relationship between robustness and consistency of connectivity measures.

## 2. Methods

### 2.1. Definition of alpha-band SNR

Alpha activity between 8 and 13 Hz is predominantly observed in occipital EEG channels. Its peak frequency and range can differ across participants. Following Nolte et al. (2008), we define an individual alpha band for each participant covering a symmetric 5 Hz range (2.5 Hz left and right of the peak) around the participant’s alpha peak frequency, where the peak is determined by maximum spectral power at electrodes O1 and O2, and is constrained to the interval [8.5, 12.5] Hz. Alpha-band signal-to-noise ratio (SNR) of an EEG sensor or reconstructed source is defined as the ratio between the spectral power at the alpha peak and the average spectral power in 2 Hz wide side bands to the left and right of the individual alpha band. Spectral power is computed using the Welch method using non-overlapping Hanning windows of 200 samples length, where we assume a sampling frequency of 100 Hz.

### 2.2. Spatio-spectral decomposition (SSD)

We apply spatio-spectral decomposition (SSD, Nikulin et al., 2011) in order to remove brain activity without strong alpha peaks. SSD seeks spatial filters w that maximize the signal power of the projected data in a frequency band of interest (here, the alpha band) while simultaneously suppressing the power in the left and right side (flanking) bands. We use the same alpha and side bands as in the definition of alpha-band SNR. Alpha band power was defined as the sum of the squared signal after 2nd order Butterworth bandpass filtering. The power in the side bands was computed analogously after application of an appropriate bandpass filter and a subsequent notch filter. Apart from these minor differences, SSD thus directly optimizes the alpha-band SNR of the projected components as defined above.

The first SSD spatial filter is given by

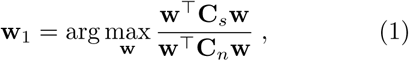

where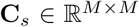 is the covariance of the sensor data filtered in the alpha band, and C_*n*_ is the covariance of the data filtered in the side bands as outlined above. A complete SSD decomposition matrix can be computed by solving a generalized eigenvalue problem (Nikulin et al., 2011).

To identify the number of SSD components, a heuristic based on the achieved alpha-band SNR of each component is employed, where only components with SNR values larger than 2 are retained for further analysis.

### 2.3. EEG source modeling

The generative model of EEG data is given by

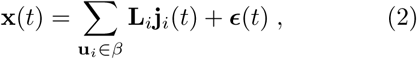

where 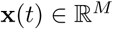 is the signal measured at *M* EEG electrodes at time *t*, 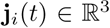 is the activity of a single source at a brain location **u**_*i*_, and where the *lead field* 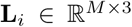 models the propagation of three orthogonal electrical point sources (dipoles) originating at **u**_*i*_ to the EEG sensors. One can rewrite the equation in matrix form as

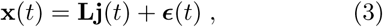

where 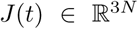 is the activity of *N* sources with 3D orientation, and 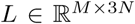 is a matrix summarizing the lead fields of *N* current sources throughout the brain. Given the geometry and electrical conductivities of the various tissues in the head, **L** can be computed; this step is called *forward modeling*. The reverse step of estimating j(*t*) given x(*t*) and **L** is called *inverse source reconstruction*.

#### 2.3.1. Forward modeling

The lead field **L** describes the physical process of neuronal current propagation from the source regions within the brain to the EEG electrodes. It can be computed based on the known geometry and electrical conductivities of the tissues in the head. Ideally, an individual geometric model should be created from a structural MRI of the participant’s own head and digitized electrode positions. However, the acquisition of individual MRI is not always possible and generally comes at a high cost. Therefore, it is common practice in EEG source analysis to use template anatomies such as the Colin27 head (a detailed MR image made of 27 scans of a single individual head, see Holmes et al., 1998) or the ICBM152 head (a non-linear average of the MR images of 152 individual heads, see Mazziotta et al., 1995; Fonov et al., 2011a).

The predominant electrical model in EEG source analysis is the boundary element method (BEM, Mosher et al., 1999). Most BEMs include realistically-shaped shells representing the brain, skull and scalp, where the electrical conductivity within each shell is assumed to be homogenous. While the BEM solution relies on numerical optimization, a quasi-analytic solution can be obtained within the same three-shell geometry using spherical harmonics expansions (SHE) of the electric lead fields (Nolte and Dassios, 2005). More accurate head models compared to the three-shell approach can be obtained using finite element based approaches (Cho et al., 2015; Vorwerk et al., 2014), however at the expense of higher computational cost.

An extension of the ICBM152 anatomy down to the neck is the so-called New York Head (Huang et al., 2015). A highly-detailed FEM solution involving six different types of tissue (scalp, skull, CSF, gray matter, white matter and air cavities) is provided by the authors for a set of 231 standardized electrode positions.

#### 2.3.2. Inverse modeling

To deal with the ambiguity of the solution of the EEG inverse problem that is caused by measuring brain activity only outside the head, it is crucial to constrain the solution to be consistent with prior domain knowledge. Common constraints include the number of sources (Scherg and von Cramon, 1986), spatial smoothness (Hämäläinen and Ilmoniemi, 1994; Pascual-Marqui, 2007), spatial sparsity (Matsuura and Okabe, 1995; Gorodnitsky et al., 1995), the combination of sparsity and smoothness (Vega-Hernández et al., 2008; Haufe et al., 2008, 2011; Sohrabpour et al., 2016), as well as constraints on the dynamics of the source time courses (Van Veen et al., 1997; Gross et al., 2001; Gramfort et al., 2013; Castaño Candamil et al., 2015).

In distributed inverse imaging, dipolar sources are modeled at many locations within the brain (in our case only in the cortical areas), and the activity at all those locations is estimated jointly. Methods that impose ℓ_2_-norm constraints on the source distribution are particularly popular, as they lead to solutions that are linear in the sensor data and therefore efficient to compute. We here consider the weighted minimum-norm estimate (WMNE Hämäläinen and Ilmoniemi, 1994), and eLORETA (Pascual-Marqui, 2007, eLORETA) as representatives for such solutions.

Another popular class of inverse methods are beamformers, which estimate brain activity separately for each source location. For each location, a beamformer finds a spatial projection of the observed signal, such that signals from that location are preserved, while contributions from all other signals contributions are maximally suppressed. The linearly constrained minimum-variance (LCMV) beamformer (Van Veen et al., 1997) does that by minimizing the variance of the filtered signal subject to a unit-gain constraint (that is, the product of filter and forward matrix at the desired location is enforced to be the identity matrix).

### 2.4. Robust connectivity estimation

The choice of the connectivity measure crucially determines not only the type of interaction that can be detected, but also whether connectivity can be reliably detected at all. It is known that various popular measures of time series interaction are not suitable for EEG-based brain connectivity analysis, as the inevitable mixing of brain sources in EEG sensors and reconstructed sources leads to excess detections of spurious connectivity based on data properties unrelated to true interaction (e.g., Schoffelen and Gross, 2009; Haufe et al., 2013). To overcome this problem, robust connectivity measures have been proposed (e.g., Nolte et al., 2004, 2008; Haufe et al., 2012, 2013; Ewald et al., 2012).

**Robustness** (w.r.t. source mixing) of a connectivity measure is defined here as the desirable property to yield zero (non-significant) results when applied to linear mixtures of independent signals.

#### 2.4.1. Functional connectivity

Functional connectivity (FC) concerns the estimation of undirected relationships between time series. Coherency is defined as the normalized version complex cross-spectrum (Nunez et al., 1997) and quantifies the linear relationship between two time series at a specific frequency. Its phase indicates the average phase difference between those series, while its absolute value (termed *coherence*) quantifies the stability of that phase delay. As such, coherence is a popular measure of functional connectivity. However, as it does not distinguish between non-zero and zero (which can be explained by source mixing even in case of independent brain sources) phase delays, it is non-robust, and may yield spurious results in practice. The imaginary part of coherency (iCOH) on the other hand is a provenly robust measure of functional connectivity as it is only non-zero for non-zero phase delays, which cannot be explained by source mixing (Nolte et al., 2004).

The empirical cross spectrum is calculated here as in Nolte et al. (2008). First, the data are divided into *K* non-overlapping segments of 2 s duration, corresponding to a frequency resolution of 0.5 Hz. Each segment is multiplied with a Hanning window before calculating the Fourier transform within the contiguous set of frequencies in the participant’s individual alpha range *F*. Denote the *k*-th segment of the *i*-th (sensor or source) time course by *x_i,k_*(*t*), and its Fourier transform by *X_i,k_*(*f*), *f* ∈ *F*. The cross-spectral matrix is defined as

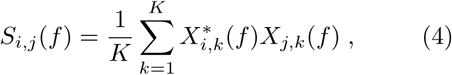

where (·)* denotes complex conjugation. Coherency between time series *x_i_*(*t*) and *x_j_*(*t*) is defined as

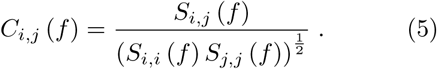

Imaginary coherence is defined as

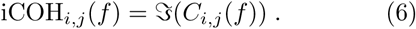

Global alpha-band imaginary coherence between two brain ROIs *p* and *q* is calculated by averaging the absolute value of the imaginary coherence across all pairs of voxels (*i, j*) within these ROIs and over all |*F*| = 11 frequency bins within the alpha range

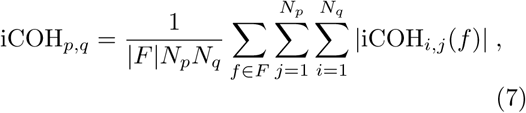

where *N_p_* and *N_q_* are the numbers of voxels inside the ROIs. Each entry of the iCOH matrix is finally divided by its standard deviation as estimated using the jackknife method to yield a standard normal distributed score

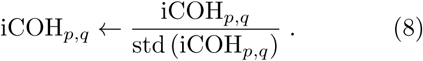

Note that, while iCOH is anti-symmetric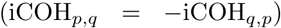, its absolute value is symmetric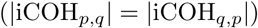.

#### 2.4.2. Effective connectivity

Effective connectivity concerns the estimation of directed interactions, in which a channel can either assume the role of the sender or the role of the receiver, or both. Granger causality (GC) (Granger, 1969) and its many variants are widely used for that purpose even though they are known to be non-robust to source mixing (Nolte et al., 2008; Haufe et al., 2013; Haufe and Ewald, 2016). The phase slope index (PSI, Nolte et al., 2008) is capable of determining band-limited effective connectivity while being robust by construction. It is therefore well suited for our investigation. PSI is based on the observation that for a constant delay between two signals, the phase of their cross-spectrum is a linear function of frequency. The sign of the slope of the phase spectrum therefore determines the lead signal. Similar to iCOH, PSI only detects non-zero delays, and is anti-symmetric. The phase-slope index between time series i and j is defined as

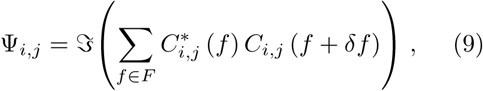

where *δf* is the frequency resolution. Global PSI between source-space ROIs *p* and *q* is obtained as

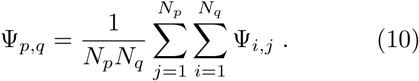

As for iCOH, we divide PSI by a jackknife estimate of its standard deviation.

### 2.5. Grand-average analysis

Grand-average SNR is obtained by averaging SNR values across all participants. Correlations between processing pipelines or participants (see the Consistency across source reconstruction pipelines and Consistency across datasets sections) are averaged across participants or pairs of participants. Denoting the standardized participant-wise iCOH or Ψ scores by *z_i_*, *i* ∈ {1,…, *N*}, grand-average scores are given by 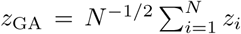. These GA z-scores can be used to test for non-zero effective connectivity. A similar test for functional connectivity would, however, require a different statistical test, as our analysis was based on absolute value of imaginary coherency, and the resulting z-scores are not symmetrically distributed around zero under the null hypothesis.

## 3. Experiments

Alpha-band oscillations constitute the strongest neural signals in the EEG. There are multiple rhythms with spectral peaks around 10 Hz that relate to different cognitive systems including the visual system (posterior alpha-rhythm) and the sensori-motor system (rolandic mu-rhythms). These oscillations are thought to represent feedback loops between the various brain structures working together to implement cognitive functions (Klimesch, 1999; Palva and Palva, 2007). While it is known that the topography of alpha oscillations is consistent with the location of the corresponding sources in occipito-parietal and central areas (Niedermeyer and da Silva, 2005; Palva and Palva, 2007), much less is known about the connectivity between different alpha sources. As each of these rhythms is strongest during inactivity of the underlying brain circuit, the resting state is ideally suited to study this question. Nolte et al. (2008) have reported directed information flow from frontal to occipital EEG sensors using PSI. In the present set of experiments, we revisit the questions using both sensor- and source-space analysis. Furthermore, we quantify the variability of source-space based results to demonstrate the uncertainty associated with anatomical interpretations of EEG source localization and connectivity analyses.

### 3.1. Data and preprocessing

For this study, we analyzed resting-state EEG data (eyes closed condition) acquired from healthy participants within two different experiments. Ethical approval was obtained for both studies.

**Fasor data (FD):** Data of N_FD_ = 30 participants (29 right-handed, one left-handed; 20 males, 9 females; age average 29.2, range 23–49) were originally recorded with 128 scalp electrodes (extended 10-20 system, 1,000 Hz sampling rate, nose reference, Easycap by Brainproducts GmbH, Munich). For the present study, we obtained a reduced dataset comprising the recordings at 64 electrodes. The recording was part of a baseline measurement embedded in an in-car EEG-study on attentional processes (Schmidt et al., 2009). Participants sat in the driver’s seat, while the car was in a parking position with the engine switched off.

**Würzburg data (WD):** Data of N_WD_ = 35 participants (28 right-handed, 7 left-handed; 13 males, 22 females; age average 25.4, range 19–35) were collected from 64 scalp electrodes (extended 10-20 system, 1,000 Hz sampling rate, right mastoid reference, Brainvision Acticap, Brainproducts GmbH, Munich) as part of a brain-computer interface study conducted in a laboratory environment. Two sessions on separate days were conducted per participant.

The length of each recording was five minutes. Data were band-stop filtered between 45 and 55 Hz, band-pass filtered between 2 and 40 Hz, and down-sampled to 100 Hz. Since source space connectivity and localization could be affected by sensor density and coverage (Hassan et al., 2014; Song et al., 2015), a subset of *M* = 49 electrodes common to both datasets was selected (see Figure 1, upper panel). Each resulting dataset consisted of an *M* × *T* multivariate time series, where *T* = 5 · 60 · 100 = 30, 000.

**Figure 1.**
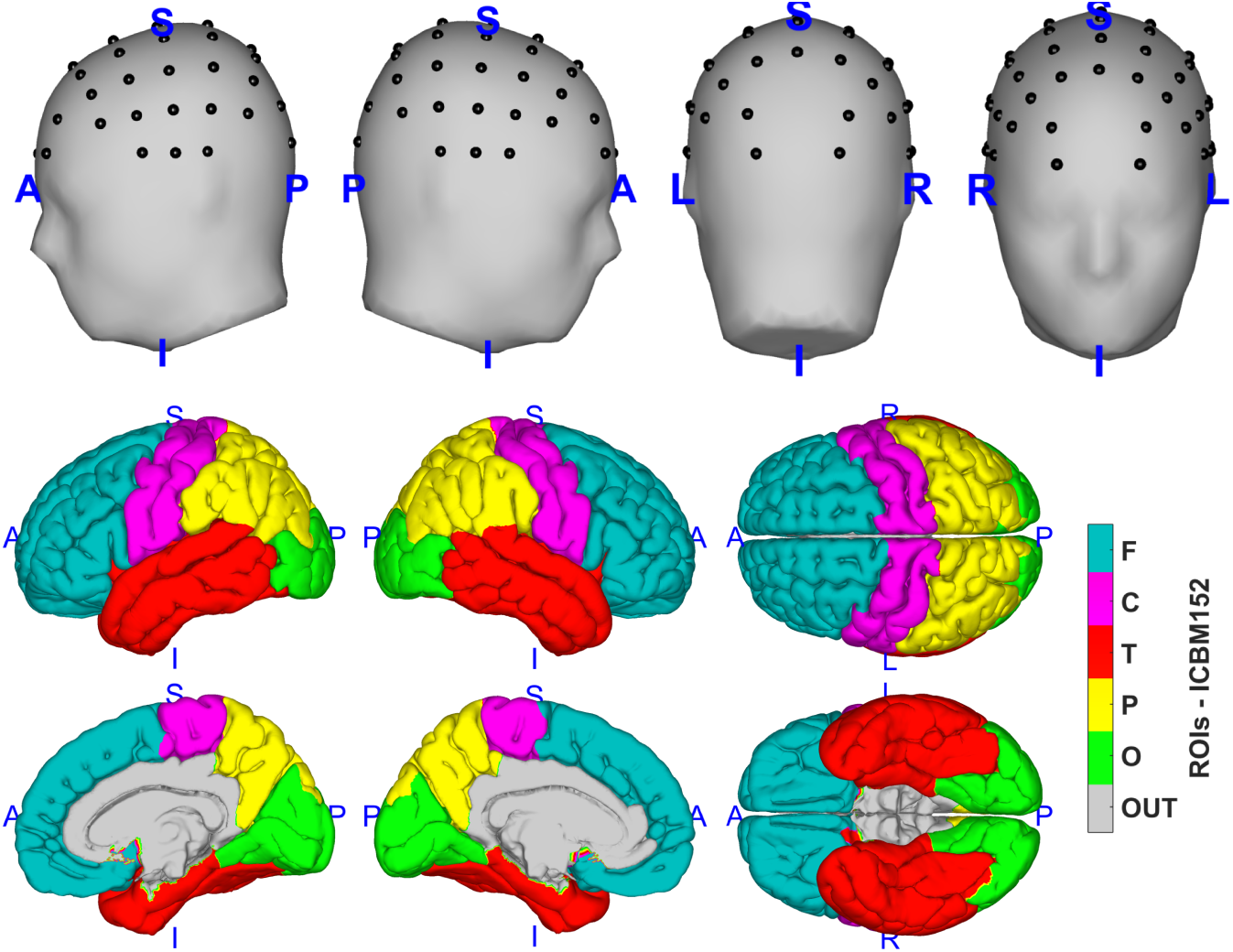
Upper Panel: Positions of the 49 EEG electrodes used in this study. Lower Panel: Regions of interest (ROI) representing the left and right frontal, parietal, temporal and sensorimotor areas, mapped onto the ICBM152 template anatomy. ROIs are defined according to the Desikan-Killiany atlas.

### 3.2. Sensor-space analysis

Nolte et al. (2008) reported global directed information flow in the alpha-band from more frontal to more occipital EEG sensors on data acquired using physically-linked mastoids as the electrical reference. We employed the identical methodology on our data to demonstrate that sensor-space results greatly depend on the electrical reference, dramatically limiting the interpretability of sensor-space connectivity analyses. On the Fasor data, we calculated grand-average sensor-space SNR and the phase-slope index for all pairs of electrodes. We then transformed the data into common average reference, and repeated the analysis. Finally, we repeated the analysis for both the Fasor and W¨urzburg data after transforming the signals to a ‘virtual’ linked-mastoids reference by subtracting the average activity from Electrodes TP9 and TP10 from each channel. As no electrodes were placed on the left and right mastoids in these studies, TP9 and TP10 were selected as the electrodes closest to the mastoids.

### 3.3. Source localization and connectivity estimation

Prior to source reconstruction, the data were transformed to common average reference. We then applied SSD to remove data components lacking a strong alpha peak. The median number of retained SSD components was 9 for the Fasor data (range 4–22), and 13 for the W¨urzburg data (range 4–25). These components were projected back to sensor space as in Haufe et al. (2014a), using the activation patterns corresponding to the SSD spatial filters (Haufe et al., 2014b).

Source reconstruction was conducted using common forward and inverse models implemented in different software packages. Specifically, we used Fieldtrip (Oostenveld et al., 2010), Brainstorm (Tadel et al., 2011), and our own Matlab-based ‘Berlin toolbox’ (Haufe and Ewald, 2016).

**Brainstorm (BS)** In Brainstorm, the EEG forward problem was solved in the ICBM152 (BS version 2015) template anatomy (Fonov et al., 2011b). Realistically-shaped surface meshes of the brain, skull and scalp were extracted from the provided template MR image using the default number of 1922 vertices per layer. The cortical surface distributed with Brainstorm was down-sampled to around 2,000 vertices. The surface was divided into ten broad regions-of-interest (ROIs) as defined by the Desikan-Killiany atlas (Desikan et al., 2006, see Table 1 for details). After excluding vertices located outside the ten ROIs (mostly voxels close to subcortical structures), *P*_BS_ = 1, 815 vertices were retained for further analysis. The forward model (lead field) from these source locations to the 49 EEG channels was calculated using three-shell BE modeling as implemented in the Open-MEEG package (Gramfort et al., 2010). The electrical conductivities used for the three compartments were σ_brain_ = 1*S*/*m*, σ_skull_ = 0.0125 *S*/*m* and σ_skin_ = 1 *S*/*m* Inverse estimation of sources was carried out using WMNE and LCMV.

**Table 1:**
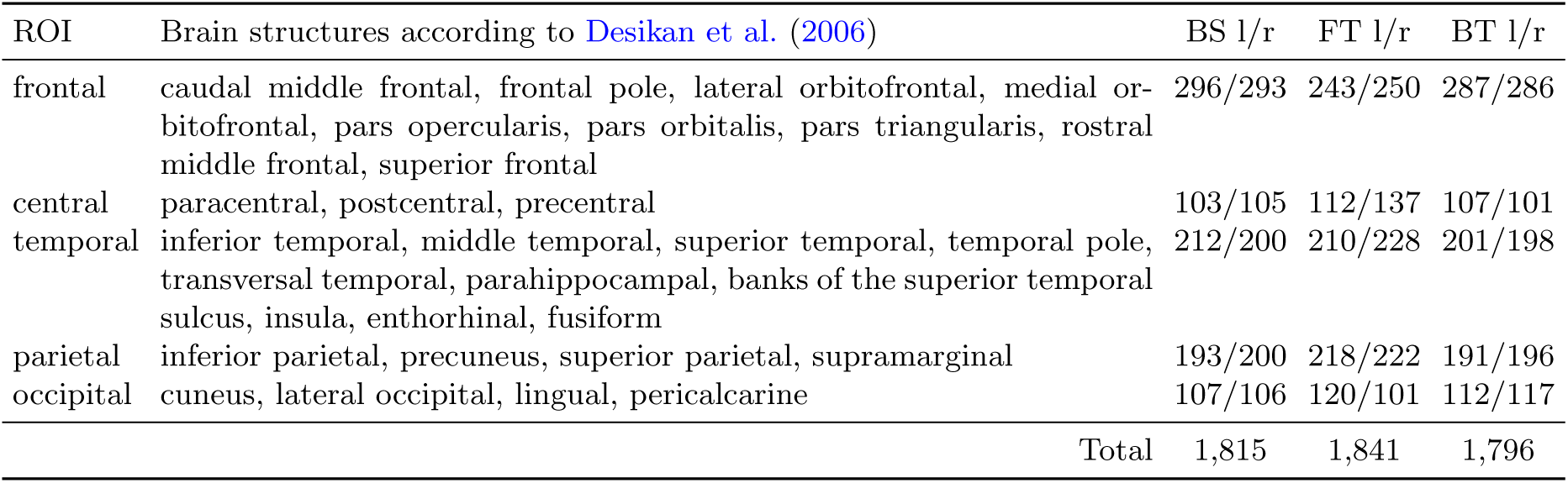
Brain structures included in the left and right frontal, central, temporal, parietal and occipital regions-of-interest (ROI), and numbers of cortical locations per ROI modeled by Brainstorm (BT), Fieldtrip (FT), and the Berlin toolbox (BT).

| ROI | Brain structures according to Desikan et al. (2006) | BS l/r | FT l/r | BT l/r |
| --- | --- | --- | --- | --- |
| frontal | caudal middle frontal, frontal pole, lateral orbitofrontal, medial orbitofrontal, pars opercularis, pars orbitalis, pars triangularis, rostral middle frontal, superior frontal | 296/293 | 243/250 | 287/286 |
| central | paracentral, postcentral, precentral | 103/105 | 112/137 | 107/101 |
| temporal | inferior temporal, middle temporal, superior temporal, temporal pole, transversal temporal, parahippocampal, banks of the superior temporal sulcus, insula, entorhinal, fusiform | 212/200 | 210/228 | 201/198 |
| parietal | inferior parietal, precuneus, superior parietal, supramarginal | 193/200 | 218/222 | 191/196 |
| occipital | cuneus, lateral occipital, lingual, pericalcarine | 107/106 | 120/101 | 112/117 |
| Total |  | 1,815 | 1,841 | 1,796 |

**Fieldtrip (FT)** Source reconstruction in Field-trip was carried out in the Colin 27 head (Holmes et al., 1998; Oostenveld et al., 2003). The cortical surface provided by FT for this anatomy was down-sampled to 2,000 vertices. In order to define ROIs on this surface, we first tesselated a Colin 27 based cortical mesh within Brainstorm using the Desikan-Killiany atlas. This mesh was then co-registered to the FT template mesh using the minimum Euclidean distance approach. Distances between vertices of the two templates were kept less than 2mm. The division of the FT template into ten ROIs left *P*_FT_ = 1, 841 vertices for further analysis. Source reconstruction was conducted using eLORETA, WMNE and LCMV. In order to circumvent a location bias toward the center of the brain, LCMV results were normalized by a noise estimate according to Equation (27) of Van Veen et al. (1997).

**Berlin toolbox (BT)** For use with the Berlin toolbox, a BE forward model of the ICBM152 v2009 anatomy (Fonov et al., 2011b) was constructed in analogy to the procedures reported for BS. To define the source space, we here however used a cortical surface provided by Freesurfer (Fischl et al., 1999; Dale et al., 1999). ROIs were defined in analogy to what is reported for FT leading to *P*_BT_ = 1, 796 source voxels. In the same geometry used for BE modeling, we also computed a forward model based on spherical harmonics expansions (SHE) of the electric lead fields (Nolte and Dassios, 2005). Finally, we also used the ‘New York Head’ representing a highly-detailed FE forward model of an extended ICBM anatomy. More details on the BE, FE and SHE modeling within the Berlin toolbox is provided in Huang et al. (2015). Sources were reconstructed using LCMV, WMNE and eLORETA.

The combination of forward and inverse models available in the various packages defined the following 14 different source reconstruction pipelines: BS-WMNE-BEM, BS-LCMV-BEM, FT-eLORETA-BEM, FT-WMNE-BEM, FT-LCMV-BEM, BT-eLORETA-BEM, BT-WMNE-BEM, BT-LCMV-BEM, BT-eLORETA-SHE, BT-WMNE-SHE, BT-LCMV-SHE, BT-eLORETA-FEM, BT-WMNE-FEM, BT-LCMV-FEM. For all pipelines, three-dimensional dipolar sources were reconstructed under the free-orientation model using the default parameters (such as regularization constants) of each package, yielding a 3*P* × *T* source times series per dataset. At source level, we then applied SSD separately to each voxel’s 3D time course. The first SSD component was retained as the dominant orientation of that voxel. In order to normalize scales across source estimation pipelines, each resulting *P* × *T* source time series was divided by its Frobenius norm. On the normalized sources of each participant, we calculated the voxel-wise alpha-band SNR as an index of the localization (LOC), as well as PSI, iCOH and COH between all pairs of ROIs.

### 3.4. Grand-average source analysis

To obtain a source-space equivalent of Figure 2, grand-average source localization maps and ROI-to-ROI functional and effective connectivity matrices were calculated from the combined data of the Fasor and Würzburg cohorts.

**Figure 2:**
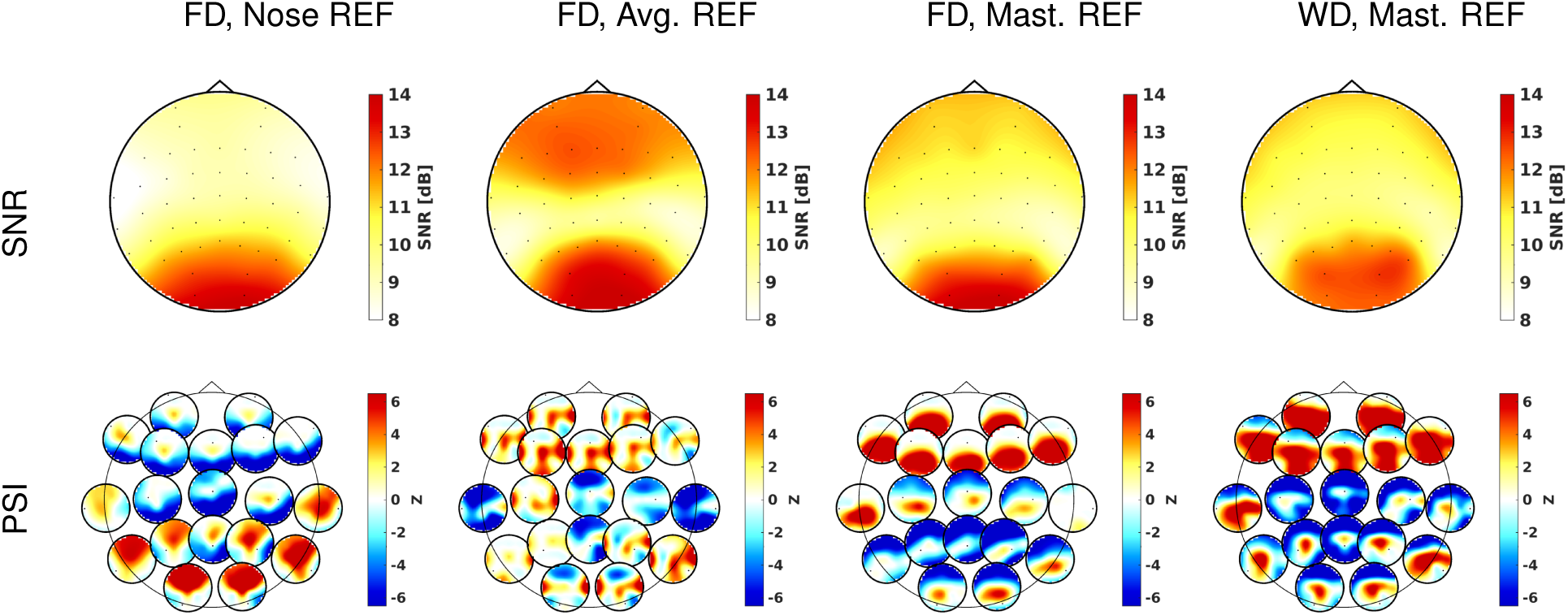
Grand-average alpha-band SNR maps calculated for 49 channels (upper panel) and effective connectivity (lower panel) computed between 19 channels according to the phase-slope index (PSI) for three different choices of the reference electrodes (nose, common average, linked-mastoids), as well as for two different datasets (FD/WD). SNR was computed per channel as the ratio of alpha-peak power and the average power in the sidebands. Effective connectivity is visualized as head-in-head plots, where red and yellow colors (*z* ≥ 2) stand for information outflow and blue and cyan colors (*z* ≤ −2) stand for information inflow. Note the similarity of the two rightmost panels with Figure 4 of Nolte et al. (2008). *The results indicate that, while sensor-space connectivity results are reproducible across datasets when the same reference electrode is used, they are substantially different for different choices of reference electrodes. They can therefore not be used to determine the locations of interaction brain sites*.

### 3.5. Consistency across source reconstruction pipelines

We quantified the consistency of source localization and connectivity results across the 14 different source reconstruction pipelines defined by the various combinations of head model, inverse solution, and implementation (software package). To this end, the source distribution of each participant was summarized in a ten-dimensional vector by averaging alpha-band SNR within each ROI. Effective and functional connectivity matrices were then stacked into 45-dimensional vectors. Consistency of the results attained by two different pipelines on the same data was assessed by means of the linear Pearson correlation between the respective localization and connectivity vectors. These correlations were computed for all pairs of pipelines, and averaged across all participants.

### 3.6. Consistency across datasets

To obtain an estimate of *between-study consistency* for each of the 14 different source reconstruction pipelines, we computed grand-average source localization and connectivity vectors separately for the Fasor and W¨urzburg data. For each pipeline, we then computed the pairwise Pearson correlation between Fasor and Würzburg results. Next, we split the Würzburg data into pairs of datasets corresponding to the first and second resting state sessions of each participant. For each participant and processing pipeline, we then computed the correlation between the results attained in the two sessions, and averaged the obtained correlations across participants to obtain estimates of inter-session or *within-participant consistency*. Finally, we computed correlations between all datasets of distinct participants separately for the Fasor and Würzburg cohorts (that is, comparisons of different session of the same participant in the Würzburg cohort were included). These correlations were averaged across all pairs of subjects to yield an estimate of for each processing pipeline and cohort.

Between-study, within-participant and between-participant consistencies were also computed on the sensor-level. To this end, ROI-based localization and ROI-to-ROI connectivity vectors were replaced by their 49- and 49·(49−1)/2 = 1, 176-dimensional sensor-space counterparts.

## 4. Results

### 4.1. Sensor-space analysis

Numerous previous studies have performed brain-connectivity estimation on the EEG sensor level, despite the fact that spurious connectivity is inevitable when applying non-robust connectivity measures to linearly mixed data (see, e.g., Haufe et al., 2013; Van de Steen et al., 2016). Here we demonstrate that, even when using robust measures, estimated connectivity patterns between EEG sensors heavily depend on the choice of the electrical reference, and are therefore hard to interpret in anatomical terms.

The results of the sensor-space analyses are depicted in Figure 2 as 2D scalp maps for SNR and as head-in-head plots for PSI-based effective connectivity. Note that SNR values are converted to a dB scale for visualization. In each head-in-head plot, each of the small scalp plots shows the estimated interaction of the corresponding electrode to the other 18 electrodes (see Nolte et al., 2008), where red and yellow colors (*z* ≥ 2) stand for information outflow and blue and cyan colors (*z* ≤ −2) stand for information inflow.

Results obtained for the approximate linked-mastoid reference are highly similar for Fasor and Würzburg data, and moreover very accurately reproduce the results reported for a exact physically-linked mastoids reference on a third dataset in Nolte et al. (2008) (Figure 4 therein). However, results obtained using different references are highly dissimilar, with nose- and linked-mastoid-referenced data even indicating reversed front-to-back interaction patterns. It can also be observed that the degree of dissimilarity is higher for effective connectivity than for SNR-based localization of alpha activity.

As the choice of an electrical reference in EEG is essentially arbitrary (even though standardizations exist, see Yao, 2001), our results emphasize that sensor data should not be interpreted in terms of the anatomical locations of interacting EEG sources and motivate the use of source reconstruction techniques to address brain connectivity questions.

### 4.2. Source localization and connectivity

Figure 3 depicts grand-average localization and connectivity results obtained for BT-eLORETA-FEM, BT-WMNE-FEM, BT-LCMV-FEM, while results of all 14 pipelines are provided in the supplement (Figures S1–3). As expected from the literature (Niedermeyer and da Silva, 2005), alpha-band sources predominantly localized in the occiptal lobes with considerable activity spreading also to temporal and parietal lobes depending on the choice of the inverse method (see top panel of Figure 3). LCMV beamforming produced SNR maps that are more focally concentrated in the occipital lobes than maps obtained using eLORETA or WMNE source imaging. The latter methods produced more blurry but highly concordant SNR distributions. The maximal alpha-band SNR achieved for WMNE and eLORETA is however higher than for LCMV (14.64 dB and 14.78 dB compared to 11.81 dB for LCMV).

Functional and effective connectivity analysis between ROIs led to more variable results than source localization, although similarities between connectivity matrices obtained on WMNE and eLORETA source estimates can be observed (lower panel of Figure 3). Connectivity analysis based on LCMV source estimates suggests left occipital as well as left and right parietal regions to be the strongest hubs of functional connectivity, while the same analysis conducted on eLORETA and WMNE estimates also designates frontal, central and temporal regions to strongly engage in FC. Left and right frontal regions are designated as net receivers of information from all other parts of the brain when working on LCMV source estimates. The picture is different when working on eLORETA and WMNE estimates, where left frontal and left and right temporal regions are determined as global senders of information.

**Figure 3:**
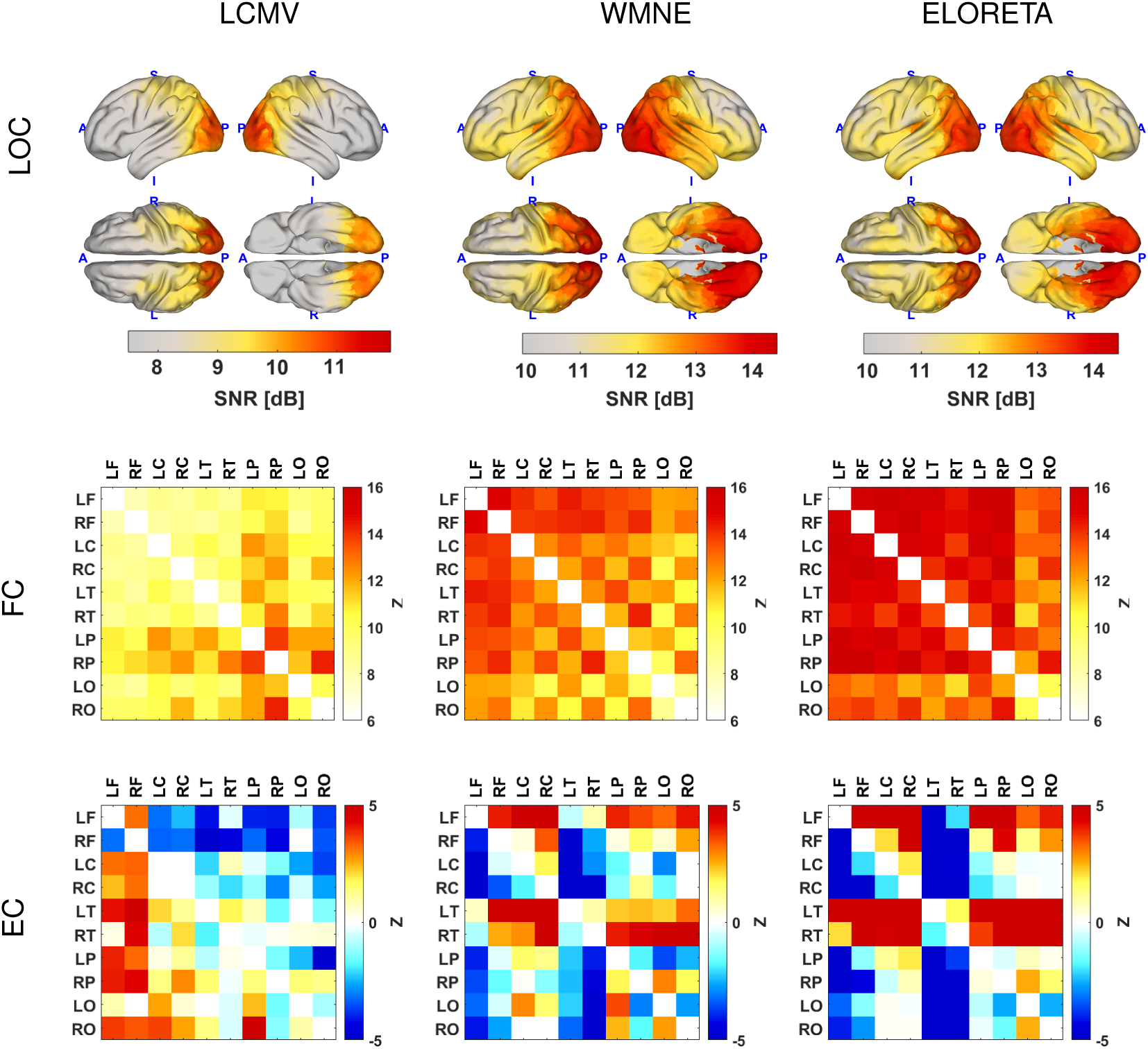
Grand-average source localization (LOC), functional (FC) and effective connectivity (EC) results obtained using finite element forward modeling and inverse source reconstruction according to linearly-constrained minimum-variance beamforming (LCMV), the weighted minimum-norm estimate (WMNE) and eLORETA as implemented within the Berlin toolbox. Upper panel: Voxel-wise relative strength (SNR) of alpha-band sources. Results are mapped onto the smoothed cortical surface of the ‘New York Head’. Center panel: Source space functional connectivity (FC) between ten ROIs as measured by the absolute value of the imaginary part of coherency. Bottom panel: Source space effective connectivity (EC) between ROIs as measured by the phase-slope index. Red and yellow colors stand for EC from rows to columns, and blue and cyan colors stand for EC from columns to rows. *The results indicate that, while sources reconstructed using different inverse methods may localize to similar brain structures, the brain interactions estimated from the reconstructed sources may differ substantially*.

### 4.3. Consistency across source reconstruction pipelines

Figure 4 depicts grand-average correlations between source localization (LOC) and functional/effective connectivity (FC/EC) results obtained using different source reconstruction pipelines on the same data. These pipelines differ w.r.t. electrical forward models (BEM, SHE, FEM), inverse models (LCMV, WMNE, eLORETA) and implementations thereof (in the BS, FT and BT packages), the latter factor also determining the template anatomy in which the source reconstruction is carried out (Colin 27 for FT, ICBM 152 for BS/BT). Source localization is found to be more consistent than source functional and effective connectivity estimation regardless of what source reconstruction parameter is varied. The average correlation across different combinations of forward models is *r* = 0.99, while values of *r* = 0.75 and *r* = 0.77 are obtained when varying inverse method and software package. Functional connectivity is more consistent than effective connectivity across all source reconstruction parameters. The average correlation observed for FC across different forward models is *r* = 0.97 compared to *r* = 0.93 for EC. For variations of the inverse method, it is *r* = 0.61 for FC compared to *r* = 0.35 for EC. Finally, the values obtained across variations of the software package/implementation are *r* = 0.70 for FC and 0.33 for EC.

**Figure 4:**
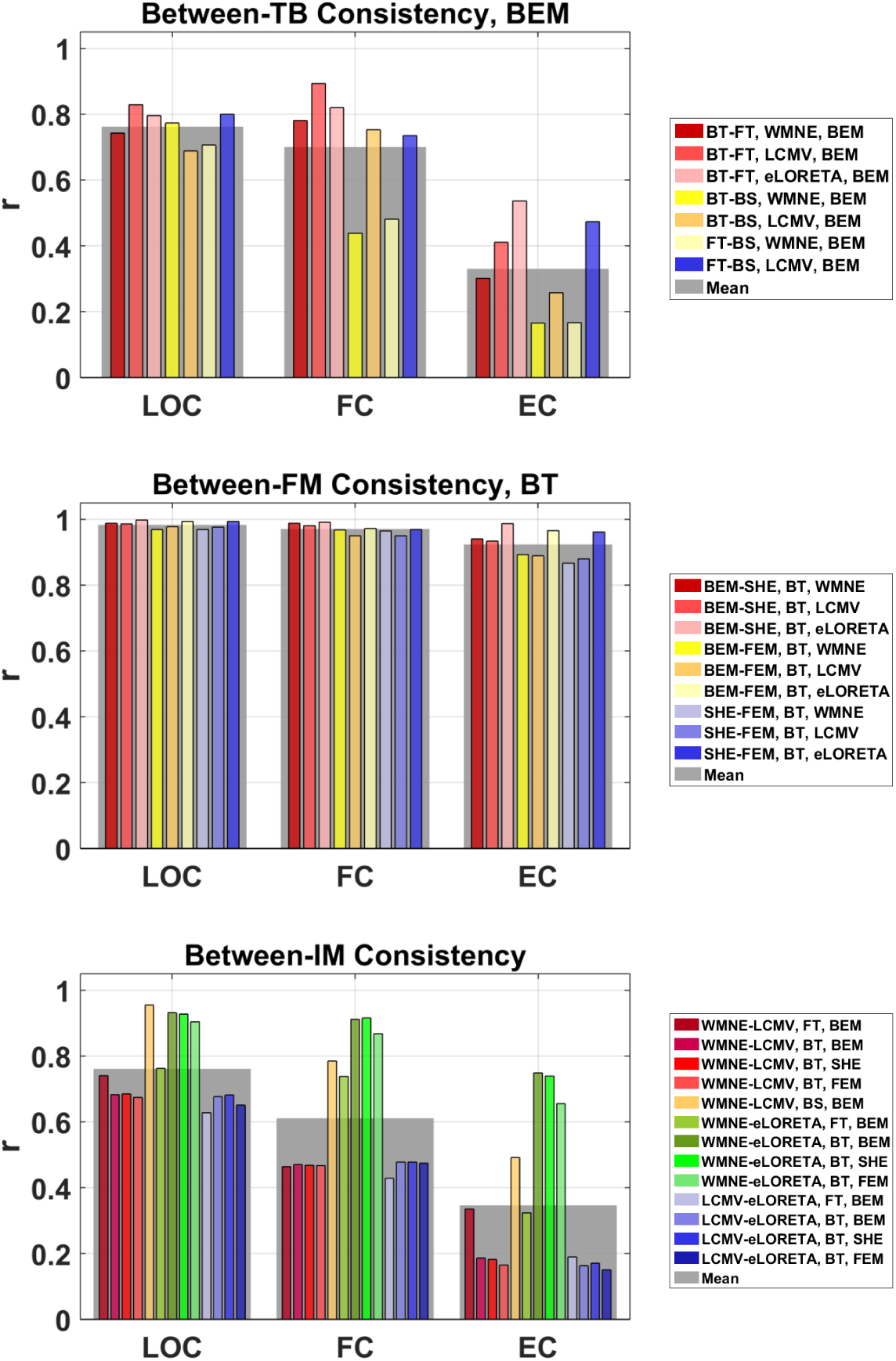
Consistency of source localization (LOC), functional connectivity (FC) and effective connectivity (EC) across source reconstruction pipelines using different forward models (FM), inverse methods (IM) and analysis toolboxes (TB). Top panel: correlation between results obtained with different software packages. BT: Berlin Toolbox, FT: FieldTrip, BS: BrainStorm. Center panel: correlation between results obtained with different electrical forward models. FEM: finite element method, BEM: boundary element method, SHE: spherical harmonics expansion. Bottom panel: correlation between results obtained with different inverse methods. WMNE: weighted minimum-norm estimate, LCMV: linearly-constrained minimum-variance beamformer. Colored bars represent the average correlation between pairs of source reconstruction pipelines. Wide grey bars indicate averages of all correlation values within each comparison subgroup. The results indicate substantial variability of source effective connectivity metrics, and to a lesser degree functional connectivity metrics and source localizations, when applied to sources reconstructed from the same data using different inverse methods or even different implementations of the same method. *The results indicate substantial variability of source effective connectivity metrics, and to a lesser degree functional connectivity metrics and source localizations, when applied to sources reconstructed from the same data using different inverse methods or even different implementations of the same method*.

Correlations between eLORETA and WMNE localizations (marked by green colors in the bottom panel of Figure 4) exceed correlations between LCMV and eLORETA (blue colors) as well as LCMV and WMNE (red colors) localizations on average by 0.20 points. For FC and EC, correlations between eLORETA and WMNE based estimates are on average 0.36 and 0.42 points higher than correlations between LCMV and eLORETA/WMNE. Not, however, that, despite this difference to WMNE and eLORETA, LCMV based estimates were highly consistent across implementations. WMNE based estimates were the least consistent across toolboxes. This result may be explained by the fact that the concept of weighted minimum-norm imaging is to reduce the influence of deep sources in the cost function, but does not precisely specify the choice of a particular weight matrix. Different implementation may therefore choose different weights (see Haufe et al., 2008, for a comparison) leading to solutions with different spatial profiles.

### 4.4. Consistency across datasets

The consistency of source localization, functional and effective connectivity estimates across studies and participants, as well as within participants is depicted in Figure 5. Highest correlations (averaged over 14 source reconstruction pipelines) are observed between grand-average results of the Fasor and Würzburg cohorts (*r* = 0.99 for LOC, *r* = 0.82 for FC and *r* = 0.48 for EC). Correlations drop when calculated on the single participant level within participants (*r* = 0.77 for LOC, *r* = 0.48 for FC and *r* = 0.30 for EC) or even across participants (WD: *r* = 0.61 for LOC, *r* = 0.32 for FC and *r* = 0.15 for EC, FD: *r* = 0.70 for LOC, *r* = 0.35 for FC and *r* = 0.15 for EC). Source localization results are most consistent between studies, within participants and between participants, followed by functional and effective connectivity estimates.

Sensor-space results were in general similarly consistent than average source-space results. Sensor maps of alpha-band activity were however less consistent between studies and participants than source localizations (*r* = 0.87 for between-study and *r* = 0.49 for between-participant consistency). Another exception is that sensor-space EC was more consistent than source-space EC between studies and within participants (*r* = 0.76 between-study and *r* = 0.48 for within-participant).

**Figure 5:**
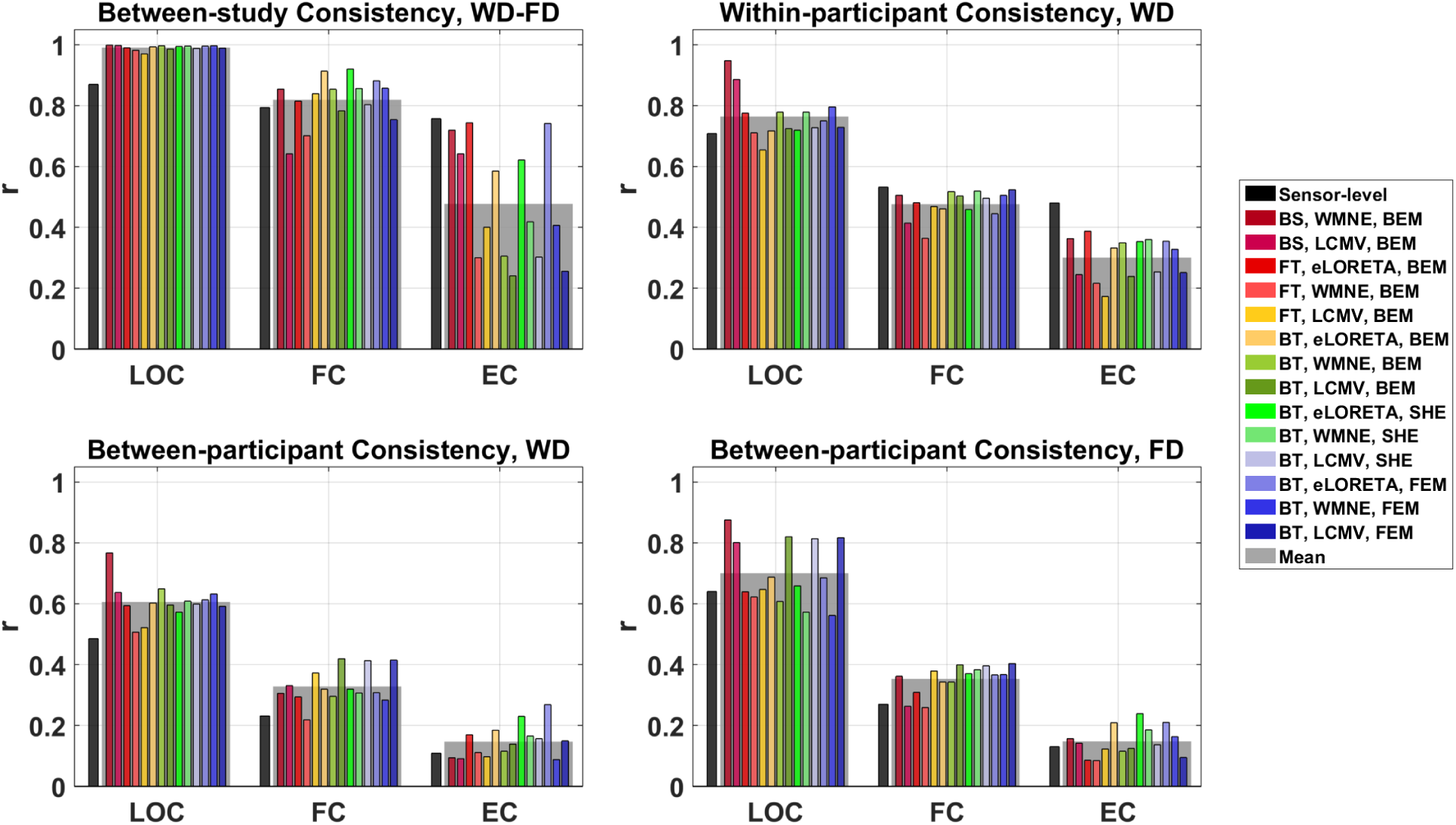
Consistency of source localization (LOC), functional connectivity (FC) and effective connectivity (EC) across 14 source reconstruction pipelines employing different forward models, inverse methods and analysis toolboxes. Top left panel: between-study (inter-dataset) consistency as measured by the correlation between grand-average results obtained for the Fasor (FD) and W¨urzburg (WD) cohorts. Top right panel: within-participant (inter-session) consistency as measured by the average correlation between results obtained from the first and second measurement session of each participant of the W¨urzburg cohort. Bottom panels: between-participant consistency as measured by the average correlation between results obtained from data of different participants within the Fasor and W¨urzburg cohorts. Colored bars represent correlations obtained for specific source reconstruction pipelines, while black bars represent analogous correlations obtained directly on sensor-space data. Wide grey bars indicate averages across all source reconstruction pipelines. *The results demonstrate that source connectivity metrics are less reproducible across participants, experimental sessions and datasets than source localizations. The lower consistency of effective compared to functional connectivity metrics suggests that resting state phenotypes are represented in different patterns of directed brain communication*.

## 5. Discussion

### 5.1. Consistency between source reconstruction parameters

The main purpose of our study was to quantify the variability of source space results that arises from the fact that EEG source reconstructions are ambiguous. To narrow down the space of possible inverse solutions, we only considered those approaches that are well established, advocated as broadly applicable, and indeed widely used. In practice, the choice of a particular source reconstruction pipeline from this pool may often just be driven by personal preference, or be based on practical concerns regarding computational complexity, availability within a certain software framework, and ease of use (e.g., automatic selection of parameters). It therefore becomes a factor that is essentially not consistent across studies and independent of the analysis goal (assuming that the choice is not based on the desired outcome). In this light, the variability of source localization, functional connectivity and effective connectivity estimates observed here across source reconstruction pipelines represents a lower bound on the uncertainty that is inherent to such estimates.

The degree of variability found in our data depends on the property of the underlying sources (location, FC or EC) that is estimated and the factor of the source reconstruction pipeline (forward model, inverse method or implementation) that is varied. The upper part of Table 2 ranks the types of analysis as well as the source reconstruction factors in terms of their observed consistency across different methods.

**Table 2:** Ranking of different analysis approaches and source reconstruction factors in terms of their consistency between methods and datasets.

|  |
| --- |
| Consistency between pipelines |
| LOC > FC > EC |
| forward model > toolbox, inverse method |
| Consistency between datasets |
| study > session > participant |
| LOC > FC > EC |

### 5.2. Consistency across different data sets

While our main results concern the variability due to different analysis pipelines applied to the same data, we also assessed the consistency of all results across different data sets. Regarding functional connectivity, we observed levels of between-study and within-participant consistency that are comparable^1^ to those reported recently in Colclough et al. (2016) for MEG data based on the imaginary part of coherency. A ranking of the data analysis and source reconstruction methods in terms of their consistency across different datasets is provided in the lower part of Table 2.

One potential explanation for the *relatively low consistency of EC and FC* across participants may be the use of resting state data. The resting state, defined as the absence of any task, leaves substantial room for participants to engage in their own thoughts. A number of distinct ‘resting state phenotypes’ has consequently been identified based on behavioral scales (Diaz et al., 2013). Given this behavioral variability, a high degree of consistency of neural metrics can not necessarily be expected.

### 5.3. Robust vs. non-robust connectivity measures

A crucial issue when estimating brain connectivity from EEG or MEG measurements is the inevitable mixing of brain sources into the measured data. The superposition of signals causes a number of connectivity metrics to yield spurious results, emphasizing once more that sensor-space connectivity analysis is inappropriate (see also Van de Steen et al., 2016). It is however worth noting that the problem of spurious connectivity also occurs at the level of source estimates as a result of the source mixing in EEG/MEG inverse solutions (e.g., Schoffelen and Gross, 2009; Haufe et al., 2013), as well as in general for any data that are superimposed by correlated noise (Vinck et al., 2015; Winkler et al., 2016).

There is currently considerable confusion regarding the question which connectivity measures can be safely applied to EEG/MEG data. One important requirement for such an application is *robustness*, defined as the property of a measure to yield asymptotically zero connectivity for linear mixtures of independent signals. Theoretical robustness has been derived for methods based on the imaginary part of coherence (Nolte et al., 2004, 2008), and on time-reversal (see below). For other approaches such as non-negative power correlations (e.g., de Pasquale et al., 2010), coherence, phase locking (Lachaux et al., 1999), and measures based on the concept of Granger causality such as the directed transfer function (Kamiński and Blinowska, 1991), partial directed coherence (Baccalá and Sameshima, 2001), spectral Wiener-Granger-causality (Bressler and Seth, 2011), and transfer entropy (Vicente et al., 2011), non-robustness can be demonstrated in simulations or using simple theoretical counterexamples (e.g., Haufe et al., 2013; Haufe and Ewald, 2016; Van de Steen et al., 2016). One such example is given by the presence of two independent non-white stationary signals (e.g., two distinct brain rhythms). Sensors or estimated sources that capture both signals with different mixing proportions will appear as interacting according to the methods listed above. Some of these approaches can however be made robust using a statistical test against time-reversed surrogate data (Haufe et al., 2013, 2012; Vinck et al., 2015; Winkler et al., 2016). Another intuitive idea to remove effects of source mixing is orthogonalization (Brookes et al., 2012; Hipp et al., 2012; Colclough et al., 2015). While respective approaches have shown encouraging results in simulations, additional theoretical analyses would be needed to rule out potentially non-robust behavior.

To avoid basing our results on spurious connectivity, we limited our analyses to robust connectivity metrics and are thus unable to report on the consistency of non-robust approaches. Yet, data provided in Colclough et al. (2016) show that non-robust connectivity metrics are typically more consistent across datasets than robust ones. This result may seem paradoxical but is rather expected, as connections determined by non-robust metrics often reflect rather simple properties of the data, for instance the strength of the sources and the corresponding mixing of the sources. Since non-robust methods are sensitive to the effects of volume conduction (source mixing) they should also reflect the spatial configuration of the sources, the latter being in fact one of the most reproducible measures in the present study. As such properties are typically quite stable across participants and repeated measurements, a high degree of consistency can be expected for non-robust connectivity measures. Robust measures, by contrast, rely on more subtle properties of the data relating to actual interactions between individual components of the superimposed signals. Such properties are harder to estimate and potentially more variable across participants and source reconstruction parameters. It is therefore important to use consistency not as the sole criterion for judging the appropriateness of a connectivity measure. Additional validation involving ground truth data (see below) is necessary to rule out that stable results are rooted in trivial biases. The same holds for all other parts of a connectivity estimation pipeline such as forward and inverse models.

### 5.4. Validation strategies

The ill-posed nature of the EEG and MEG inverse problems entails substantial uncertainty not only in the locations of underlying brain sources, but also in the results of all analyses conducted on reconstructed sources such as source connectivity analyses and the subsequent estimation of network properties of connectivity graphs. The accumulation of errors in such complex analysis chains calls for thorough validation. We postulate that validation efforts should aim to estimate not only biases but also variances contributing to the overall error. Here we contribute to these efforts by estimating the variance of EEG source analyses that is introduced by the choice of source reconstruction parameters.

Future studies should reproduce our work using simulated data, where the known ground truth can be used to assess the correctness of the results. Current simulation studies on EEG source reconstruction are limited by focusing on localization accuracy as the sole performance criterion; thus not taking into account the correct recovery of time series dynamics that would be required for subsequent connectivity analysis. Conversely, many existing studies of time series connectivity neglect the residual source mixing that is present in reconstructed brain sources. In order to overcome these limitation, we have suggested that future studies should jointly benchmark and compare entire source connectivity estimation pipelines (Haufe and Ewald, 2016).

In addition to simulations, it is further important to also analyze real-world data with known ground truth. Such data may be obtained from artificial ‘phantom’ heads. Alternatively, EEG recordings obtained either during deep brain stimulation or during concurrent recording of invasive electrophysiology could be used (e.g., Papadopoulou et al., 2015). Finally, it is important to also intensify the theoretical study of connectivity measures and source imaging procedures.

In the light of the currently insufficient understanding of what source imaging methods are best suited for reconstructing brain sources with (specific types of) dependencies, we suggest meanwhile to use more than one inverse modeling approach for the estimation of source connectivity patterns based on EEG data. Ideally these approaches should not be based on the same mathematical framework in order to avoid consistency of the results due to the possible common algorithmic assumptions. A convergence of the results from alternative methods is a promising sign and a basis for further functional interpretation of the findings. If, however, multiple algorithms strongly indicate different results, e.g.: opposite flow of information or missing/present interactions between the structures of interest – a further analysis and validation of the data is strongly encouraged.

### 5.5. Limitations and future work

While EEG-based brain connectivity analysis can be performed in numerous different ways, we had to limit our approaches to the variation of only a few processing parameters, while all other parameters had to be fixed to values that we deemed reasonable and representative of pipelines that are typically used in practice. While we have outlined the necessity of using robust connectivity measures for EEG and MEG data above, many other of our choices may be challenged. In particular, the number of electrodes may be regarded too low to perform source localization. Unfortunately, no high-density dataset was available for the present study. However, we did not observe substantially different results (e.g., higher between-methods consistency) for slightly larger sets of 57 (FD) and 61 (WD) electrodes (see Supplementary Figures S4– 9). Similarly, it may be argued that a larger number of ROIs would have been more representative of state-of-the-art studies. However, to simplify the estimation tasks, we deliberately chose a small number of ROIs. A finer tessellation of the cortical surface into 60 ROIs consequently led to less between-method consistency (see Figures S4–8 and S10). One may further argue that, while the use of SSD dimensionality reduction has been shown to improve source localization accuracy in simulations (Haufe et al., 2014a), SSD is not yet an integral part of many source connectivity estimation pipelines. We therefore conducted additional experiments, in which we omitted the application of SSD in sensor space, while we replaced SSD by principal component analysis (PCA) when determining the dominant direction of alpha-band activity within each source voxel. These analyses demonstrate a general decrease in the consistency between different inverse solutions when using PCA instead of SSD (Figures S4–8), which is likely caused by SSD’s ability to extract oscillatory signal components with higher SNR than PCA.

A further limitation of our study may be seen in the use of resting state data. Future studies should investigate whether a higher degree of consistency can be obtained by contrasting different experimental conditions. Furthermore, a comparison of our results to the MEG case would be of considerable interest. Finally, an open question to be addressed in future work is to what extent the observed variability between inverse methods and even implementations thereof is due to incongruent choices of higher-level parameters such as regularization constants and depth-compensation strategies.

## 6. Conclusion

Our data show that EEG-based source localizations as well as source-leakage-corrected functional and effective source connectivity estimates display a considerable dependency on the choice of source reconstruction parameters among common options. This variability reflects uncertainty in the results, and should be discussed when reporting findings based on source reconstructions in future studies. In the absence of clear evidence for the superiority of a particular methodology, researchers may want to report source space analyses based on more than one EEG source reconstruction pipeline.

## Acknowledgements

SH was supported by a Marie Curie International Outgoing Fellowship (grant No. PIOF-GA-2013-625991) within the 7th European Community Framework Programme. VVN acknowledges a support by the Russian Academic Excellence Project ‘5-100’. We thank Guido Nolte for contributing Matlab code.

## Footnotes

1 Note that the quantities reported in Colclough et al. (2016) are Fisher z-transformed correlations ρ = atanh(r).

